# Quantifying the unquantifiable: why Hymenoptera – not Coleoptera – is the most speciose animal order

**DOI:** 10.1101/274431

**Authors:** Andrew A. Forbes, Robin K. Bagley, Marc A. Beer, Alaine C. Hippee, Heather A. Widmayer

## Abstract

**Background:** We challenge the oft-repeated claim that the beetles (Coleoptera) are the most species-rich order of animals. Instead, we assert that another order of insects, the Hymenoptera, are more speciose, due in large part to the massively diverse but relatively poorly known parasitoid wasps. The idea that the beetles have more species than other orders is primarily based on their respective collection histories and the relative availability of taxonomic resources, which both disfavor parasitoid wasps. Though it is unreasonable to directly compare numbers of described species in each order, the ecology of parasitic wasps – specifically, their intimate interactions with their hosts – allows for estimation of relative richness. We present a simple logical model that shows how the specialization of many parasitic wasps on their hosts suggests few scenarios in which there would be more beetle species than parasitic wasp species. We couple this model with an accounting of what we call the “genus-specific parasitoid-host ratio” from four well-studied genera of insect hosts, a metric by which to generate extremely conservative estimates of the average number of parasitic wasp species attacking a given beetle or other insect host species. Synthesis of our model with data from real host systems suggests that the Hymenoptera may have 2.5 - 3.2× more species than the Coleoptera. While there are more described species of beetles than all other animals, the Hymenoptera are almost certainly the larger order.

---

> “…if the micro-hymenopterists would get off their lazy asses and start describing species, there would be more micro-Hymenoptera than there are Coleoptera.”
>
> — – Terry Erwin (in [1])

The beetles (order Coleoptera), have historically [2–4] and contemporaneously [5–10] been described as the most speciose order of animals on Earth. The great diversity of beetles was sufficiently established by the middle of last century such that J.B.S. Haldane (possibly apocryphall^1^) quipped that an intelligent creator of life must have had “…an inordinate fondness for beetles” [11]. However, what evidence underlies the claim that the Coleoptera are more species-rich than the other insect orders? Certainly, more species of beetles (>350,000) have been *described* than any other order of animal, insect or otherwise [12], but does this reflect their actual diversity relative to other insects?

Why are beetles thought to be so diverse in the first place? In part, historical biases in beetle collecting and an associated accumulation of taxonomic resources for the Coleoptera may have had an outsized influence on our perception of diversity. In the mid-to-late 1800s, beetles were prized among insects for their collectability. Many landed gentlemenincluding, notably, Charles Darwincollected beetles for sport and would make a great show of comparing the sizes of their respective collections [13,14]. This preconception was then reinforced by studies that extrapolated from specific, targeted collections of insect diversity that focused on beetles. Of these, perhaps the highest in profile was a study conducted by Terry Erwin. Erwin [15] used an insecticide to fog the canopies of 19 individual *Luehea seemannii* trees in a Panamanian rainforest and then collected and identified the insect species that fell out of those trees. After having identified the proportion of the beetle species that were apparently host-specific to *L. seemannii* (163 of 955), he estimated that there might be as many as 12.2 million beetle species in the tropics. Similar studies seeking to estimate global insect diversity have also tended to emphasize beetles (e.g., [16,17]).

Nevertheless, some previous work has challenged the canon, with various authors suggesting – though never quite insisting – that the Hymenoptera may be more speciose than the Coleoptera [18–21]. The premise behind this suggestion is that most of the larvae of the Parasitica (one of the two infraorders of apocritan Hymenoptera; the other is the Aculeata, which includes ants, bees, and wasps), are obligate parasites of insect and other arthropod hosts that feed on the host’s tissue until the host dies (≈ “parasitoids”). Why is this parasitic life history relevant to the Hymenoptera’s proportional contribution to insect diversity? Simply put, species of parasitoid Hymenoptera (including the Parasitica, as well as some other groups such as the Orussidae and some Chrysidoidea) attack all orders of insects as well as some non-insect arthropods [22–24], and, reciprocally, most holometabolous insect species are attacked by at least one – and often many more than one – species of hymenopteran parasitoid [25,26]. For instance, Hawkins and Lawton [27] examined parasitoid communities associated with 158 genera of British insects across five different orders, and found that parasitoid species richness ranged from 2.64 – 9.40 per host species across different host insect orders.

If parasitoid wasps are ubiquitous and most hosts are attacked by many different species, why is there any debate at all about the Hymenoptera being more diverse than other orders? One reason may be that estimates of the regional and global species-richness of parasitoid wasps remain elusive. Their small size and a relative paucity of taxonomic resources have left the parasitoid Hymenoptera relatively under-described compared to other insect orders [20,28]. As a consequence, when collection-based estimates of regional insect diversity have been attempted, they have often excluded all but the largest and easiest-to identify families of parasitic Hymenoptera (e.g., [29–31]; though see [32,33]).

A second reason for uncertainty regarding the species richness of the parasitoid Hymenoptera is that their host ranges are often unknown. While it may be true that most insects harbor many parasitoid species, the question remains whether these parasitoid communities are exclusively composed of oligophagous or polyphagous wasps that attack many hosts, or if instead the average insect host tends to have some number of specialist wasps among its many predators (**Figure 1**). Only in the latter case would one be able to confidently assert that the Hymenoptera is the largest of the insect orders.

**Figure 1.**
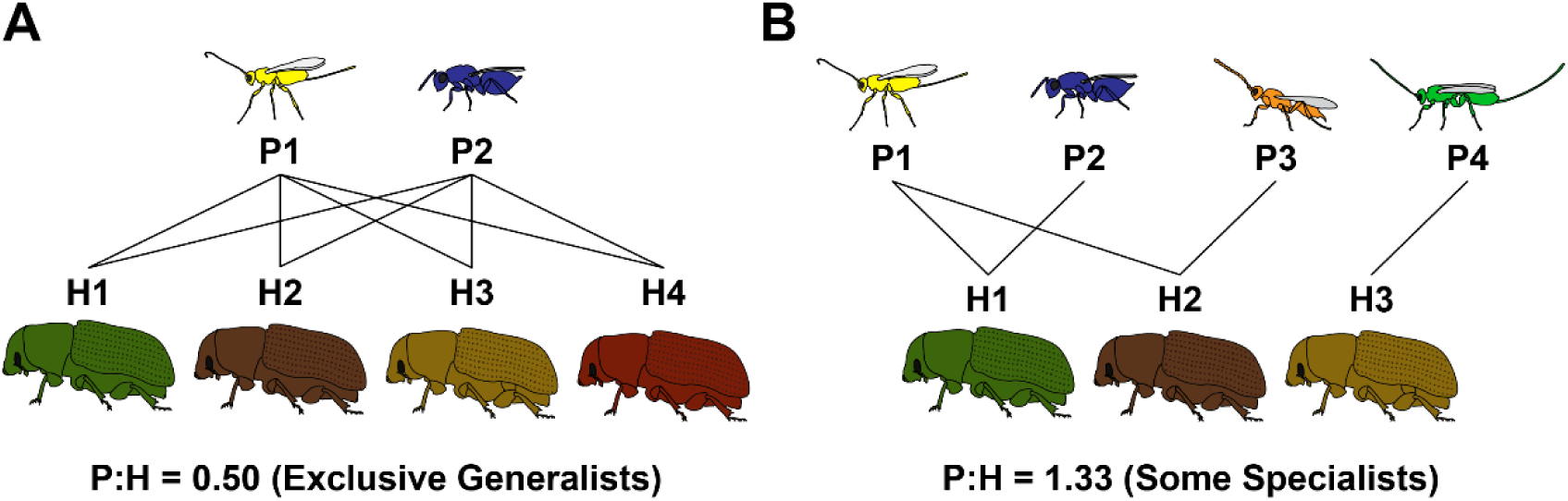
An illustration of how uncertainty about specialist vs. generalist behaviours might lead to misleading conclusions about parasitoid species richness. In panel A, each host species (differently colored beetles) is attacked by two parasitoids. However, because all parasitoids attack all four beetles the overall species richness of hosts exceeds that of the parasitoids (i.e., P:H < 1). In panel B, while some hosts have only one parasitoid, overall parasitoid richness exceeds host richness (P:H > 1) because some parasitoids are more specialized.

How then to approach this question without asking the micro-hymenopterists (and the coleopterists, dipterists, lepidopterists, etc.) to hurry up and describe all of the world’s insect species? We suggest two complementary approaches: 1) mathematically describing the values of parasitoid-to-host (“P:H”) ratios that would support – or contradict – the notion that the Hymenoptera is the most speciose insect order and 2) tabulating – wherever possible – actual P:H ratios for various genera of host insects.

## What parasitoid-to-host ratios would suggest that the Hymenoptera are more species-rich than other insect orders?

For the Hymenoptera to be the largest order of insects, the global ratio of wasp parasitoids to hosts (P:H) need not – in fact – equal or exceed 1.0. Indeed, a global P:H of 1.0 (i.e., an average of one unique hymenopteran parasitoid species for each other insect species) would mean that parasitoids account for a full half of all insects. Instead, P:H ratios need only reach values such that the Hymenoptera are more species-rich than the next largest order (which, for the sake of argument, we will assume is the Coleoptera). Here, we work towards finding parameters that describe that space. First, it will be true that:

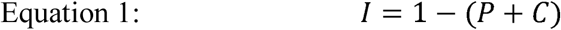

Where *P* is the proportion of all insect species that are parasitoid Hymenoptera, *C* is the proportion of insects that are Coleoptera, and *I* is the remaining proportion of insect species (**Figure 2 *A***). Note that *I* includes the non-parasitoid Hymenoptera while both *I* and *P* exclude the many Hymenoptera that are parasitic on other parasitoids (“hyperparasitoids”).

**Figure 2.**
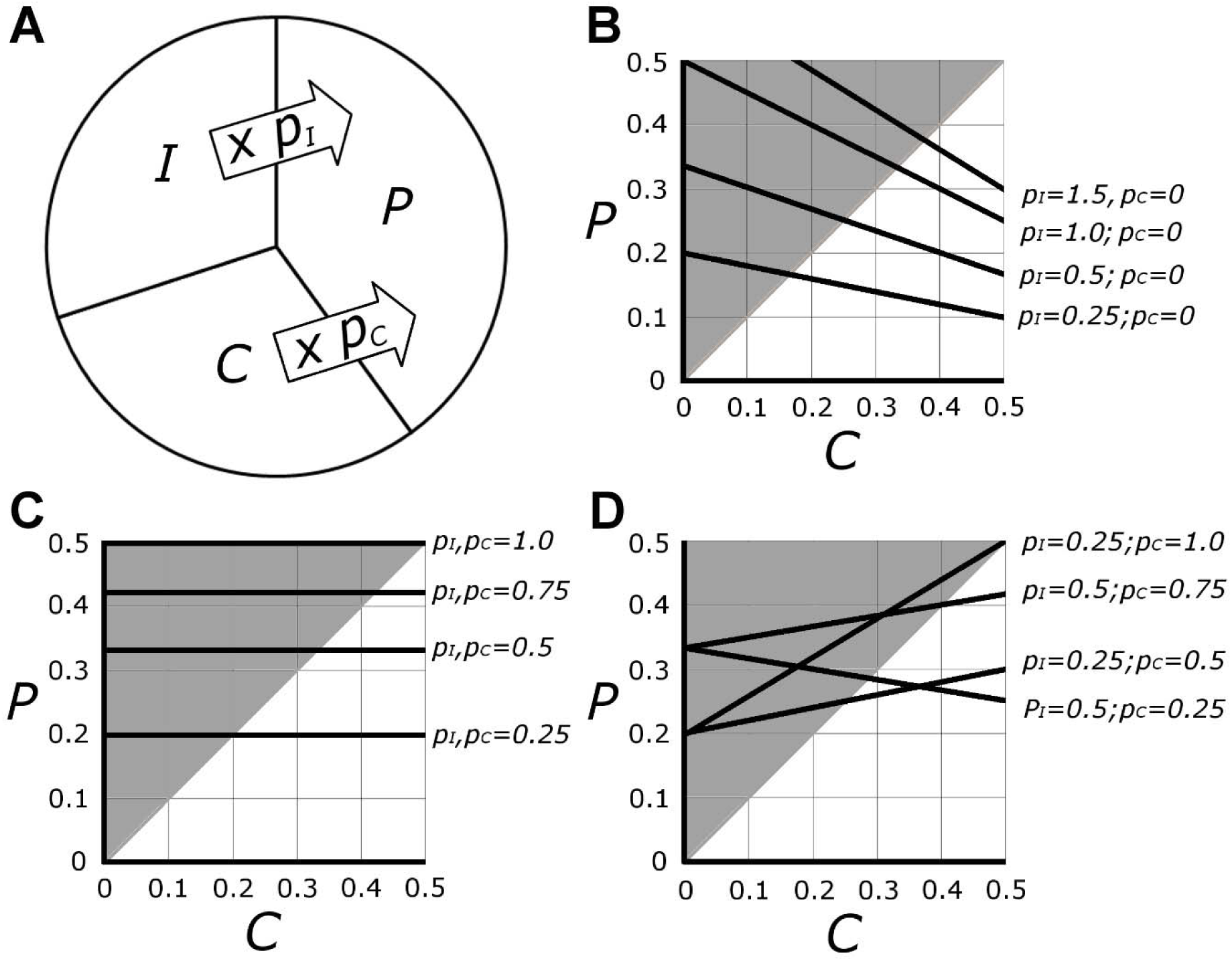
Representations of the space where the number of parasitoid wasp species would outnumber the Coleoptera, given different parasitoid-to-host ratios for coleopteran hosts and for other insect hosts. A) Pictorial representation of the model, wherein the total number of parasitoid species (*P*) will be the sum of the number of species of Coleoptera (*C*) and of other insects (*I*), each first multiplied by their respective overall parasitoid-to-host ratio (*p*_*C*_ or *p*_*I*_); B) Black lines show results of the model for four different values of *p*_*I*_and with *p*_*C*_ held at zero (i.e., when the average coleopteran has no specialist parasitoids). Where black lines overlap with gray shaded areas represents space where *P* > *C;* C) Results of four different scenarios in which *p*_*C*_ and *p*_*I*_are equal; D) Some additional combinations of *p*_*C*_ and *p*_*I*_. Though both axes could continue to 1.0, some high values of *P* and *C* are not mathematically possible or biologically likely, and at *P* or *C* values above 0.5 the question about relative species-richness becomes moot.

Additionally, because of the intimate relationship between parasitoids and their hosts, we can describe the proportion of species that are parasitoid Hymenoptera using the following expression:

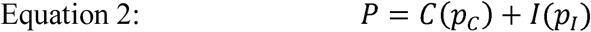

where *p*_*C*_ and *p*_*I*_represent the mean P:H ratios for all coleopterans and all non-coleopterans, respectively. The true values of *p*_*C*_ and *p*_*I*_ are unknowable, but can be estimated (see next section), and their use in this way allows for exploration of the ranges of P:H ratios that would result in different relative numbers of Hymenoptera and Coleoptera. Equation 2 again excludes hyperparasitoids, as well as parasitoids of non-insect arthropods, which makes *P* a conservative estimate of the proportion of insect species that are parasitoids.

Given these two relationships, we can substitute Eq.1 into Eq. 2:

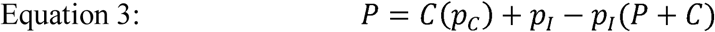

Equation 3 allows us to find the values of *p*_*C*_ and *p*_*I*_ that result in a *P* > *C* or vice versa. As shown in **Figure 2**, the space where *P* > *C* includes a substantial area where *p*_*C*_ or *p*_*I*_(or both) can be < 1. For instance, if the Coleoptera make up 25% of all insects, as suggested by many contemporary authors [17,34], a *p*_*C*_ of only 0.25 (or one species-specialist parasitoid for every four beetle species), coupled with a *p*_*I*_ of 0.50, results in *P* = *C* (and the many tens of thousands of non-parasitoid Hymenoptera will then tip the scale in their favor). Even if the Coleoptera amount to 40% of the insects, which reflects the percentage of currently-described insect species that are beetles, there will be more parasitoid Hymenoptera than beetles if *p*_*C*_ and *p*_*I*_ are equal to or in excess of 0.67 (two specialist parasitoid species for every three host species).

Another way to explore the values of *p*_*C*_ and *p*_*I*_ at which *P* will be greater than *C* is find the moments when the two will be equal. If we substitute *C* for *P* into Eq.3, we get:

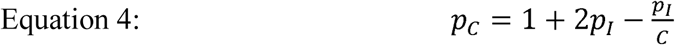

We can then plot *p*_*C*_ vs *p*_*I*_ for values of *C* between 0 and 0.5 (**Figure 3**). Here, each line represents moments when *P* = *C*, such that the area above and to the right of each line represents values of *p*_*C*_ and *p*_*I*_ that result in a *P* > *C*. Here again, *p*_*C*_ and *p*_*I*_ need not be particularly large for the parasitoid Hymenoptera to exceed the species richness of the Coleoptera. For instance, if one quarter of all insects are beetles, *p*_*C*_ and *p*_*I*_ need only exceed 0.4 (the equivalent of two parasitoid species for every five host species).

**Figure 3.**
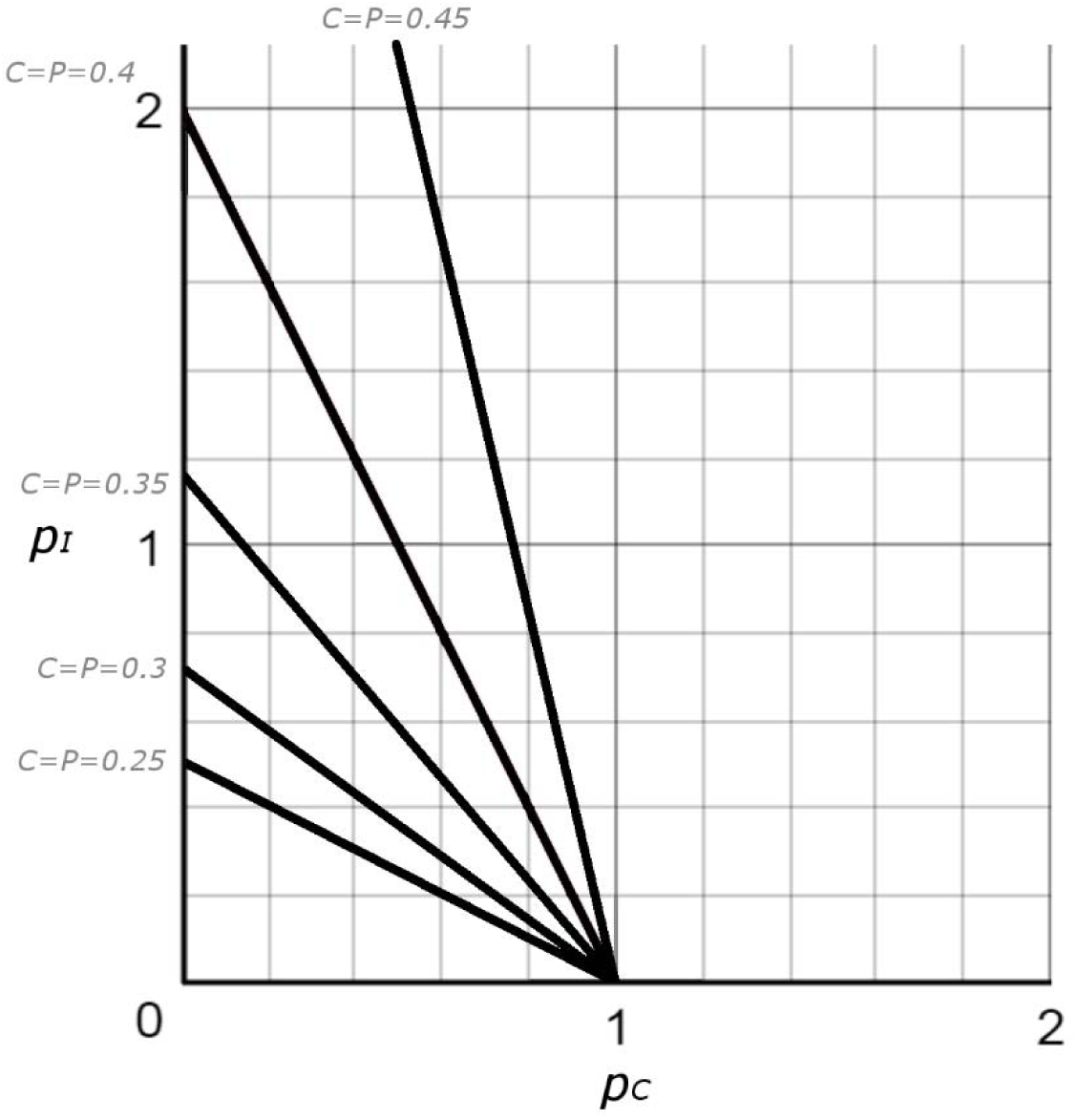
Plot based on Equation 4, with five representations of moments when *C* and *P* are equal proportions (solid black lines). *p*_*I*_ = overall P:H ratio for non-coleopteran insect hosts; *p*_*C*_ = overall P:H ratio for the Coleoptera. Space above and to the right of each line represents values of *p*_*C*_ and *p*_*I*_ where *P* > *C*, while space below and to the left of each line represents values where *C* > *P*.

## What do actual P:H ratios look like in nature?

The next question becomes: can we estimate parasitoid: host ratios (e.g., *p*_*C*_, *p*_*I*_) for different host insects? Quantifying global P:H ratios for entire insect orders is as unapproachable as the task of counting all of the living insect species: not only are most Hymenoptera undescribed, host records for described species are often incomplete, such that multiplying each host species by its supposed number of specialist parasitoids may often inadvertently include parasitoids that share hosts (**Figure 4**). While this is problematic, recognition of the problem helps present paths forward. For indeed, *some* host-parasitoid systems are exceedingly well studied and well-understood, such that we can be reasonably confident about the completeness of the host records of at least some parasitoids. With this information, we can calculate a metric that we call the genus-specialist parasitoid:host ratio. This metric interrogates all members of a host insect genus in the same geographic region and identifies all of the parasitoids known to attack only members of that genus (the “genus-specialist” parasitoids). Because this P:H ratio ignores all parasitoids known to attack any extra-generic host – as well as those whose host range is unknown or has been incompletely studied – it is therefore an extremely conservative estimate of the overall P:H ratio for an insect genus.

**Figure 4.**
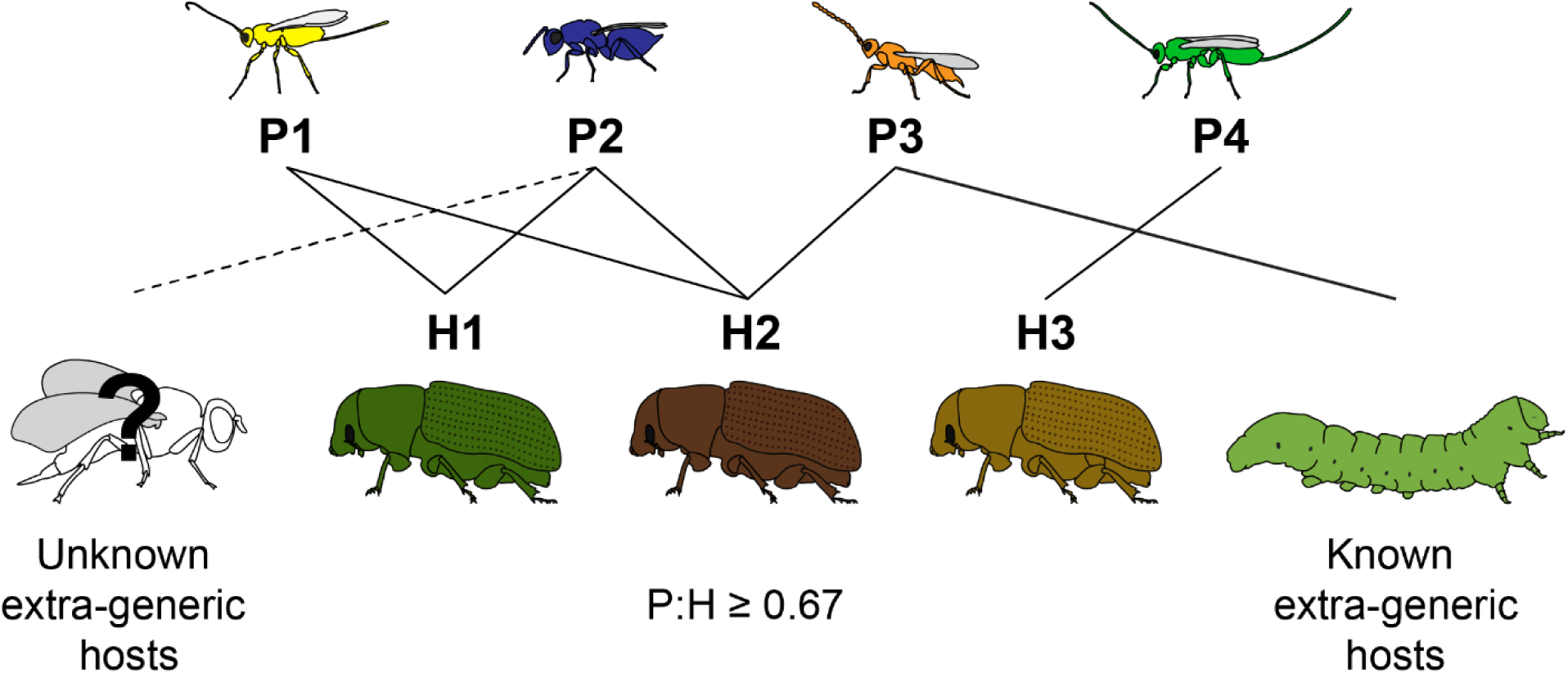
Known genus-specialist parasitoids can be used to calculate a minimum P:H ratio for an insect host genus. The focal beetle genus H (three species) has four known parasitoids, P1-P4. P1 and P4 are relatively well-studied, and known to be genus-specialists, attacking only hosts in this beetle genus. P3 has some known extra-generic hosts, while the host range of P2 is poorly studied and unknown extra-generic hosts may exist. For the purposes of estimating a genus-specialist P:H, one would therefore use only P1 and P4, such that a minimum P:H for this beetle genus would be 2/3, or 0.67. Note that if the total number and identities of extra-generic hosts were known for P2 and P3, a “true” P:H for the genus could be calculated (see **Synthesis**, below).

Below, we present four case studies, representing host-parasitoid systems with records sufficiently complete to allow for calculation of genus-specialist parasitoid:host ratios. For each system, we focus on a single host genus in North America. We restricted geography so that parasitoid numbers would not be inflated by large biogeographic differences between hosts in their parasitoid assemblages. North America was chosen because sampling is relatively strong, and several robust resources exist for Nearctic parasitoids (e.g., [24,35,36]).

For each system, we searched for all literature that mentioned the name of the host genus (or historical synonyms) and either “parasite” or “parasitoid” and compiled a database of records, performing reticulated searches on each parasitoid species name as it was added to the database in order to determine known parasitoids host ranges. From among all parasitoid records, we classified parasitoids as “genus-specialists” if they had only ever been reared from hosts in this same genus. We then split these “genus-specialists” into two groups: those for which an argument can be made that they do not have unknown extra-generic hosts, and those that were “possible genus-specialists” but for which records were less complete. Non-hymenopteran parasitoids (e.g., Tachinidae) were excluded, but in any case were only present for two of the four hosts we examined (*Malacosoma* and *Neodiprion*), and generally do not have the taxonomically cosmopolitan host ranges of the hymenopteran parasitoids. For cases where host genera were found on multiple continents, only host species in North America were included in the study, and to be conservative, a parasitoid was still considered “generalist” if it occurred on an extra-generic host species outside of North America. Introduced host species were noted but not counted in host lists, as they do not represent long-term host-parasite relationships. Introduced parasitoid species were included in generalist lists, regardless of whether they were specialists on that genus in North America or elsewhere. We describe each system below and refer the reader to Supplemental Materials for species lists, specialist / generalist classifications, and citations. A summary of data across the four genera can be found in **Table 1**.

**Table 1.**
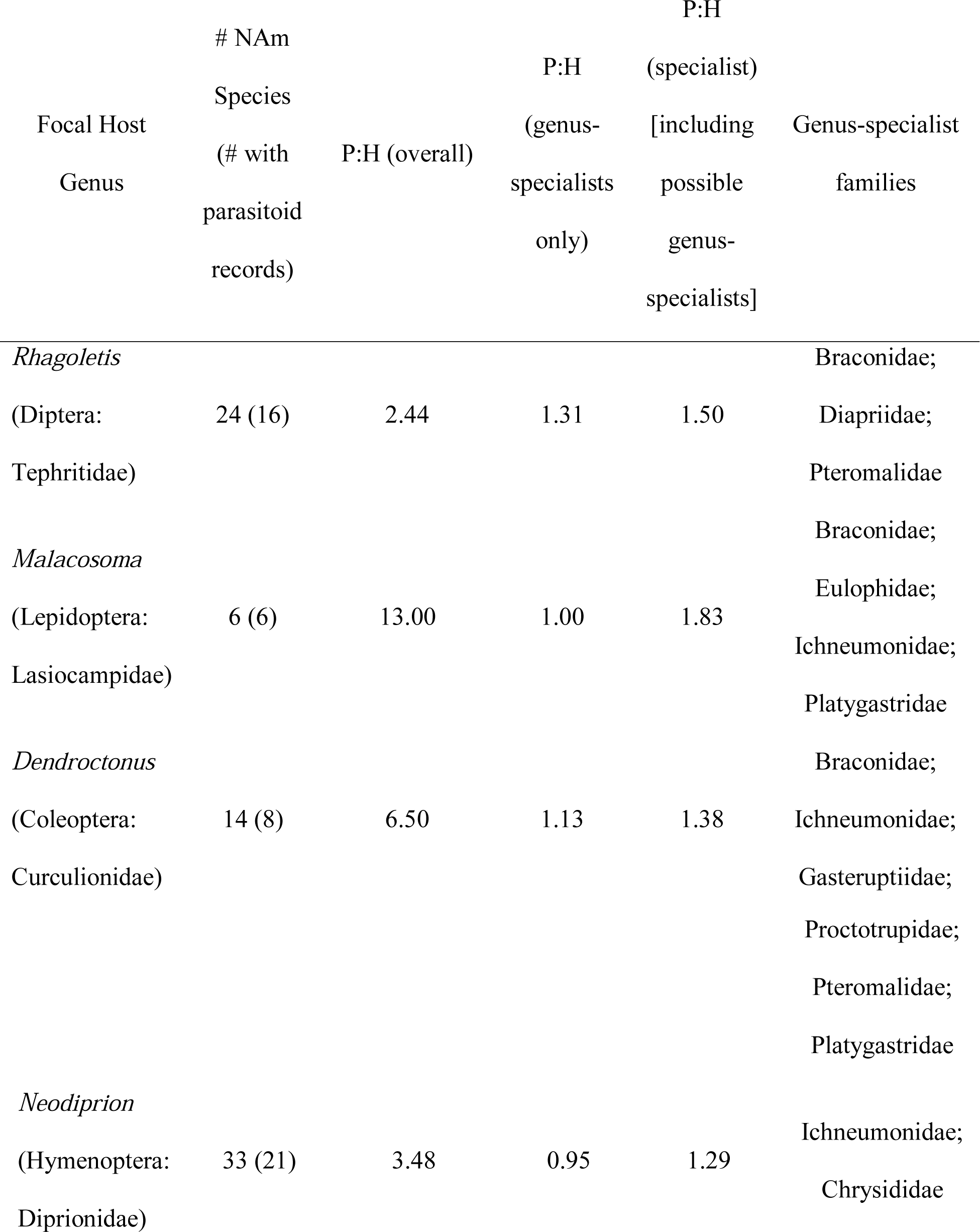
Summary of estimates of parasitoid to host (P:H) ratios for four host insect genera. Shown for each host genus are: the total number of North American (NAm) species, as well as the number with parasitoid records; the overall P:H, which includes generalist species; the genus-specialist P:H; and the genus-specialist P:H when “possible genus-specialists” were included. Parasitoid families that were among each group of genus-specialists are also listed.

### System 1: *Rhagoletis* (Diptera: Tephritidae)

Many North American *Rhagoletis* flies are pests of agriculturally-important fruits. Eggs are deposited in ripening fruits by the female fly, and larvae develop through several instars while feeding on fruit pulp [37]. For most species, larvae then exit the fruit and pupate in the soil. Parasitoids are known from egg, larval and pupal stages of many *Rhagoletis* species. Several studies have described the parasitoid communities associated with *Rhagoletis* agricultural pest species (e.g., [37–41]), though records of parasitoids of non-pest species also exist (e.g., [42– 44]). Moreover, many of the associated parasitoid species are well-studied in their own right, with robust records of their biology, ecology, and host-ranges [40,45–47].

Of the 24 species of North American *Rhagoletis* flies, 16 have a published record of parasitoid associations. Across these 16 flies, we found records of 39 parasitoid species, among which 24 “genus-specialists” have been described only from North American *Rhagoletis* and no other insect host (**Supplemental Table 1**). Of these, we set aside three “possible” genus-specialist species that did not have a strong collection record and for which host records may possibly be incomplete. The remaining set of genus-specialists included 14 braconids (genera *Diachasma, Diachasmimorpha, Utetes*, and *Opius*), six diapriids (genus *Coptera*), and a pteromalid (genus *Halticoptera*). The genus-specialist P:H ratio for *Rhagoletis* is therefore either 1.31 (21/16), or 1.50 (24/16), depending on whether “possible genus-specialists” are included. An extra-conservative P:H ratio might also include the eight *Rhagoletis* hosts that have no record of parasitoids (P:H = 21/24 = 0.88), though this almost certainly ignores some number of unknown genus-specialist parasitoids.

Some of the 15 “generalist” parasitoids of *Rhagoletis* have been reared from a diverse set of extra-generic hosts, but in some cases only from one other fruit-infesting tephritid (e.g., *Phygadeuon epochrae* and *Coptera evansi*, both of which have only been reared from *Rhagoletis* and from *Epochra canadensis* [Diptera: Tephritidae]). These 15 “generalists” are listed in **Supplemental Table 1**.

### System 2: *Malacosoma* (Lepidoptera: Lasiocampidae)

The tent caterpillars (genus *Malacosoma*) are shelter building, cooperatively-foraging moths that damage both coniferous and deciduous trees across at least 10 families. Most species use >1 host tree genus, though some (e.g., *Malacosoma constrictum*; *Malacosoma tigris*) are more specialized [48]. There are six North American species of *Malacosoma*, some with overlapping geographic distributions [48]. Female moths lay eggs in a mass wrapped around a branch of the host tree. Larvae of most species (*M. disstria* is an exception) live colonially inside “tents” made of spun silk and make regular excursions to feed on host leaves. The caterpillar stage is eaten by birds, mammals and several insect predators, but the most taxonomically diverse natural enemies are the parasitoids [48]. Of these, approximately one third are Dipteran (family Tachinidae), while the remaining two thirds are Hymenopteran parasitoids. Parasitoids attack all immature life stages, but most appear to emerge during the pre-pupal or pupal stage. Parasitoids of the North American tent caterpillars have been well documented, and often in the context of other available forest caterpillar hosts, such that it is reasonable to assert that some parasitoid species are *Malacosoma*-specific (e.g., [49–51]).

All six of the North American *Malacosoma* species have at least one known parasitoid association, and we compiled a total of 78 different parasitoid species across all hosts (**Supplemental Table 2**). Of these, eleven had only been reared from *Malacosoma*. Five of these eleven species we assigned to the “possible genus-specialists” category, as they had not been assigned a specific name (which makes it hard to determine whether other hosts exist), or because they had only been reared a single time from the host. The remaining six “genus-specialists,” were from four different hymenopteran families. The genus-specialist P:H ratio for *Malacosoma* is therefore between 1.00 and 1.83.

**Table 2.**
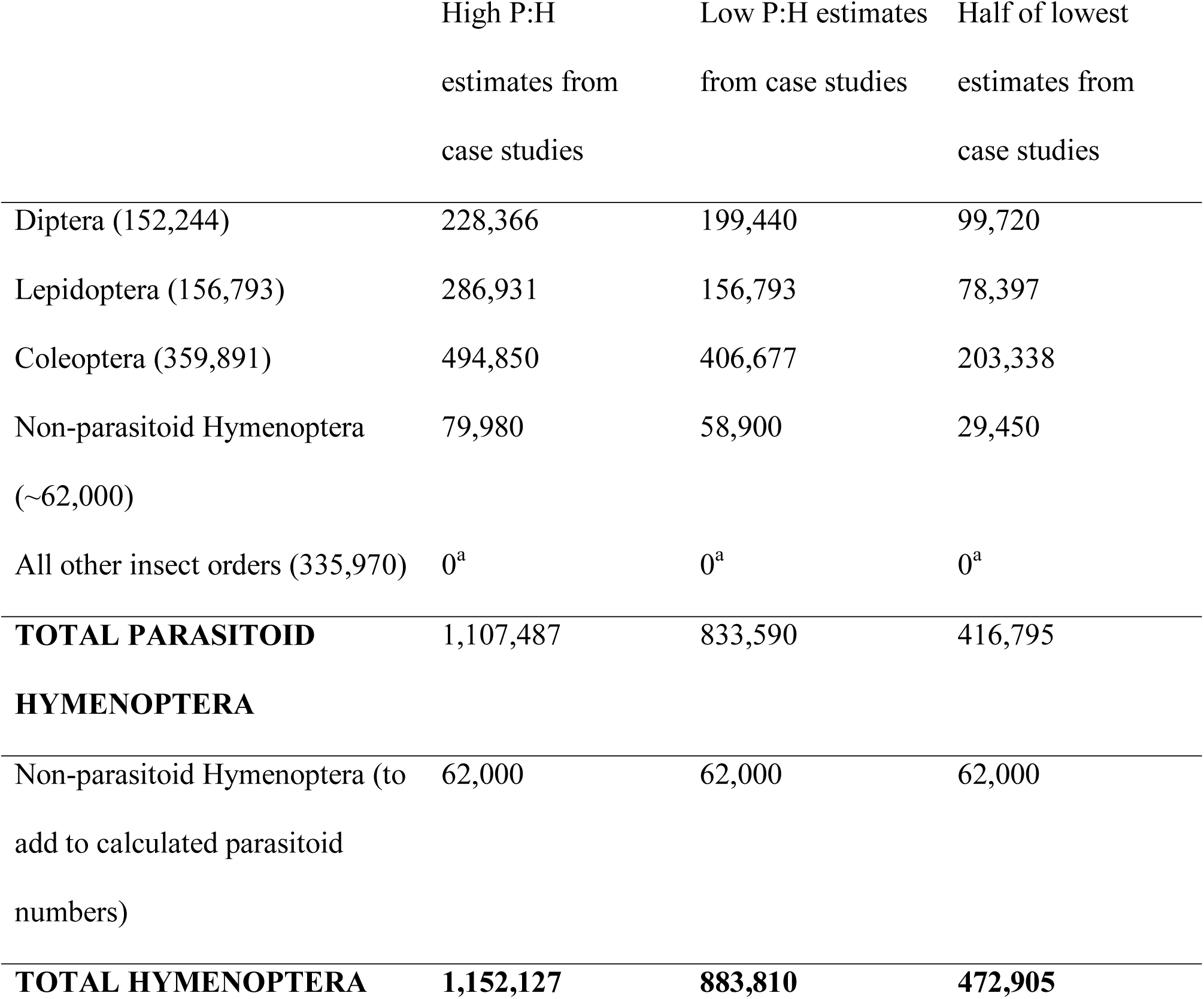
Calculations of hymenopteran species richness, given numbers of described insect species in other orders and P:H ratios estimated in this paper. Combining conservative P:H ratio estimates from four case studies with numbers of described species in the four largest insect orders [28,70] offers an idea of how species richness of the Hymenoptera may compare with that of other orders. ^a^Parastioids attack hosts in all other insect orders, but these are omitted as we did not estimate P:H ratios for any hosts in these orders. Total numbers therefore exclude large numbers of hymenopteran species.

*Malacosoma* have many more “generalists” than *Rhagoletis*: 68 species have been reared from both *Malacosoma* and at least one other extra-genetic host (**Supplementary Table 2**). Many of these appear to be specific to Lepidopteran hosts.

### System 3: *Dendroctonus* (Coleoptera: Curculionidae)

Approximately 14 species of *Dendroctonus* bark beetles are found in North America [52]. *Dendroctonus* are specific to conifers in family Pinaceae, and can be highly destructive to their host trees. Female beetles construct nuptial chambers in trees where they mate with males and then deposit eggs in tunnels in the phloem. Larvae feed on phloem and outer bark and leave the tree only after pupation and adult emergence [52]. Most species are tree genus- or species-specific.

Parasitoids have been described for eight of the 14 North American *Dendroctonus* species, though for two of these (*D. adjunctus* and *D. murryanae*) only one or two parasitoid species are known. The total list of *Dendroctonus*-associated parasitoids is long, but the records are also often problematic, as *Dendroctonus* share their habitat with several other genera of bark beetles, which may or may not be attacked by the same parasitoids. In many studies, parasitoids are listed as “associates” of either *Dendroctonus*, or of one of the other species, or of both, but this does not always necessarily mean that a parasitoid attacks that beetle [53–55]. We have here again tried to be conservative, though in one case (*Meterorus hypophloei*) we have ignored a claim of “association” with *Ips* beetles [56] as it did not seem to be well justified and other authors describe *M. hypophloei* as a *Dendroctonus frontalis* specialist [55,57]. In total, we found nine *Dendroctonus* genus-specialists, two possible genus-specialists, and 48 “generalists” (**Supplemental Table 3**). The genus-specific P:H ratio for *Dendroctonus* is therefore between 1.13 and 1.38.

### System 4: *Neodiprion* (Hyemenoptera: Diprionidae)

*Neodiprion* is a Holarctic genus of pine-feeding sawflies specializing on conifers in the family Pinaceae [58]. These sawflies have close, life-long associations with their tree hosts. The short-lived, non-feeding adults mate on the host plant shortly after eclosion, after which the females deposit their eggs into pockets cut within the host needles. The larvae hatch and feed externally on the host needles throughout development, and then spin cocoons on or directly beneath the host [59–61]. Many species also have highly specialized feeding habits, and feed on a single or small handful of host-plant species in the genus *Pinus*. Since many of the ∼33 *Neodiprion* species native to North America are considered economic pests [62], considerable effort has gone into describing their natural history and exploring potential methods to control *Neodiprion* outbreaks.

Despite the wealth of natural history information, compiling a list of parasitoids attacking *Neodiprion* is complicated by a history of accidental and intentional introductions. In addition to the native species, the European pine sawfly, *Neodiprion sertifer*, and three species from the closely related genera *Diprion* and *Gilpinia* were introduced in the past ∼150 years and have spread across the United States and Canada [63–66]. In an attempt to control these invasive pests, several parasitoids have been introduced, and now attack both native and invasive diprionids [67–69].

We found 20 genus-specialist parasitoid species associated with the 21 species of North American *Neodiprion* for which parasitoid records exist. An additional seven parasitoids were classified as “possible” genus-specialists. The genus-specific P:H ratio for *Neodiprion* is therefore between 0.95 and 1.29. An additional 51 species had been reared from both *Neodiprion* and an extra-generic host, with nine introduced parasitoids. We also compiled a list of 14 introduced parasitoids, nine hyperparasitoids, and 28 tachinid (Diptera) parasitoids of *Neodiprion* (**Supplemental Table 4**), but these were not included in any analyses.

### Synthesis

Upon considering our model together with actual estimates of P:H ratios from natural host systems (**Table 1**), there appear to be few conditions under which the Hymenoptera would not be the largest order of insects. If, for instance, the P:H ratios for *Rhagoletis, Malacosoma, Dendroctonus*, and *Neodiprion* are at all representative of other hosts in those respective orders, and we use them to calculate relative species richness based on recent counts of only the *described* species in each order [70], the Hymenoptera exceed the Coleoptera by 2.5-3.2 times (**Table 2**). Recall that these calculations ignore all hyperparasitoids, and also omit parasitoids of other insect orders (e.g., Hemiptera, Orthoptera) and of non-insect arthropods. Even if we use half of the lowest P:H ratio estimate for each of the four largest orders, the Hymenoptera would outnumber the Coleoptera by more than 1.3 times.

Note that P:H ratios might be measured more accurately and / or calculated in different ways, most of which we would expect to increase the estimates of P:H reported here. For instance, rather than ignoring all of the so-called “generalist” parasitoids, one could identify those for which host ranges are known (e.g., **Figure 4**), divide each by the total number of host genera attacked, and add that fraction to the numerator of the P:H ratio for the focal host genus. As one example, the “generalist” parasitoids *Phygadeuon epochrae* and *Coptera evansi* both attack only *Rhagoletis* flies and the currant fly *Epochra canadensis*. These would each add an additional 0.5 to the other 24 “genus-specialist” parasitoids of *Rhagoletis*, giving a revised P:H of 1.56. For *Malacosoma, Dendroctonus*, and *Neodiprion*, which all have many “generalist” parasitoids with host ranges that include only a few other extra-generic hosts in the same respective family, such additions should increase P:H ratio estimates by a considerable margin.

Another way to calculate P:H would be to focus not on a host genus but on hosts sharing the same habitat. For instance, *Dendroctonus* bark beetles share their habitat niche with several other species of beetle, and many of their parasitoids are “specialists” in the sense that they attack more than one bark beetle, but all within the same tree habitat [55]. One could, therefore, calculate a P:H where H is the number of potential beetle host species in the habitat, and P is the number of “habitat-specialist” parasitoid species (those that attack one or more of the hosts in that habitat and no other hosts in other habitats).

Our analyses largely ignore the increasingly common finding that many apparently polyphagous insects – both herbivores and parasitoids – show evidence of additional host-associated genetic structure that might, if considered here as distinct lineages, change P:H ratios (e.g., [71–75]). Indeed, all four of our focal host genera have named subspecies or show evidence for host-associated, reproductively-isolated lineages [52,76–78]. Though we chose to “lump” subspecies and other reproductively isolated lineages together for this analysis, it is interesting to consider how a detailed study of genetic diversity and reproductive isolation among a host genus and all of its associated parasitoids might change P:H ratios. Studies of the flies in the *Rhagoletis pomonella* species complex and three of their associated parasitoids suggest that where additional host-associated lineages are found in a phytophagous insect, this cryptic diversity may be multiplied many times over in its specialist parasitoid community [46,79]. If broadly true, this implies that genus-specific P:H ratios may often be much higher than we report here.

One sensible criticism will surely be: to what extent are the P:H ratios for these four genera reflective of global P:H ratios for their respective orders (Coleoptera, Lepidoptera, Diptera, and the non-parasitoid Hymenoptera)? Surely some insect genera escape parasitism, and perhaps the examples chosen here simply have exceptionally large, or unusually specialized, parasitoid communities. As to the former, it may be that such escape artists exist, but they also may be relatively rare. After all, there are parasitoids that attack aquatic insects [80,81], that parasitize insects in Arctic communities (e.g., [82]), and even those that dig down into soils to unearth and oviposit into pupae [45]. The list of potential hosts for parasitoids also extends to many non-insect arthropods, including spiders, mites, and nematodes [83,84]. As to the four example genera being representative of overly large parasitoid communities, all of their “overall” P:H numbers (**Table 1**) are actually below the means found for their respective orders in an extensive study of parasitoid communities in Britain [27], suggesting that these communities are of average, or slightly below-average, size.

### Concluding Thoughts

While it may indeed be premature to claim that the Hymenoptera is the largest order of insects based solely on our data, many other studies offer support for the same conclusion. In fact, the preponderance of evidence suggests that the common wisdom about the Coleoptera being the most speciose is the more dubious claim. Studies of insect diversity that reduce taxonomic biases have found the Hymenoptera to be the most species-rich in both temperate [32] and tropical [33] forests, as well as in other habits (e.g., [85,86]). In addition, a mass-barcoding study of Canadian insects found both Hymenoptera and Diptera more diverse than Coleoptera [87]. Moreover, other historically-accepted ideas about diversity of parasitoid hymenopterans have recently been questioned, including the apparent myth that parasitoids are one of only a few groups whose diversity decreases towards the tropics [88,89]. In any case, we hope this commentary results in a redoubled effort to understand and describe the ecology and natural histories of parasitoid wasps, including host ranges and cryptic host-associated diversity, such that estimates of P:H can be made for additional host genera. We also hope to see similar efforts in other animal groups that may harbor great diversity but for which far too little is known about host ranges, such as particularly speciose orders of mites and nematodes (e.g., [90,91]. In other words, and to again quote Erwin [15], we hope that “…someone will challenge these figures with more data.”

## Authors’ Contributions

AAF conceived of the study. All authors helped formulate a framework for addressing the questions in the paper, developed the logical model, and collected and analyzed data from the four host genera. AAF and RKB wrote the manuscript. All authors gave final approval for publication.

## Acknowledgements

We thank Isaac Winkler, Anna Ward, Eric Tvedte, Miles Zhang, Glen Hood, and Matt Yoder for their thoughtful discussions and comments on this manuscript.

## Footnotes

1 Whether or not Haldane ever actually said it exactly in this way is unresolved [11]. This phrase does not occur in any of Haldane’s writing, but he does write that “The Creator would appear as endowed with a passion for stars, on the one hand, and for beetles on the other.” [92]

