## Supplementary Materials for "Quantifying the unquantifiable: why Hymenoptera – not Coleoptera – is the most speciose animal order"

Jiménez Quiroz, E., Sánchez Escudero, J., Equihua Martínez, A., Montiel, M., Tuliaseso, J., and Valdez Carrasco, J. 2008. Distribución, abundancia y parasitismo de *Ooencyrtus kuvanae* (Howard) (Hymenoptera: Encyrtidae) parasitoide de los huevos de *Malacosoma incurvum* Hy. Edwards (Lepidoptera: Lasiocampidae) en Xochimilco, DF (No. TESIS.). Colegio de Postgraduados, Campus Montecillo, Campus Montecillo, Postgrado de Fitosanidad, Entomología y Acarología.

Juliano, S. A., and Borowicz, V. A. 1987. Parasitism of a frugivorous fly, *Rhagoletis cornivora*, by the wasp *Opius richmondi*: relationships to fruit and host density. Canadian journal of zoology 65:1326-1330.

Kapler, J. E., and Benjamin, D. M. 1960. The biology and ecology of the red-pine sawfly in Wisconsin. Forest Science 6(3).

Katovich, S. A., McCullough, D. G., and Haack, R. A. 1995. Yellowheaded spruce sawfly – It’s ecology and management. United States Department of Agriculture Forest Service General Technical Report NC-179:1-24.

Knerer, G., and Wilkinson, R. C. 1990. The biology of *Neodiprion pratti* (Dyar) (Hym., Diprionidae), a winter sawfly in West Florida. Journal of Applied Entomology 109:448-456.

Kraemer, M. E., and Coppel, H. C. 1978. The parasitoids of the European pine sawfly *Neodiprion sertifer* (Geoffroy) (Hymenoptera: Diprionidae), in Wisconsin, with keys to adults and larval remains. Wisconsin Academy of Sciences, Arts and Letters 66:91-112.

Krombein, K.V., Hurd, P.D., Smith, D.R. and Burks, B.D., 1979. Catalog of Hymenoptera in America north of Mexico. Smithsonian Institution Press. Washington, D.C.

Krugner, R., Daane, K. M., Lawson, A. B., and Yokota, G. Y. 2005. Biology of *Macrocentrus iridescens* (Hymenoptera: Braconidae): a parasitoid of the obliquebanded leafroller (Lepidoptera: Tortricidae). Environmental entomology 34:336-343.

Kulhavy, D., and Miller, M. C. 1989. Potential for biological control of *Dendroctonus* and *Ips* bark beetles. Stephen A. Austin State University Faculty Publications. 215.

Kulman, H. M. 1965. Natural control of the eastern tent caterpillar and notes on its status as a forest pest. Journal of Economic Entomology 58:66-70.

Langor, D. W. 1991. Arthropods and nematodes co-occurring with the eastern larch beetle, Dendroctonus simplex [Col.: Scolytidae], in Newfoundland. Entomophaga 36:303-313.

Langor, D. W., and Raske, A. G. 1987. Reproduction and development of the eastern larch beetle, *Dendroctonus simplex* LeConte (Coleoptera: Scolytidae), in Newfoundland. The Canadian Entomologist 119:985-992.

Langston, R. L. 1957. A synopsis of hymenopterous parasites of *Malacosoma* in California. University of California Publications in Entomology 14:1-50.

Lee, J. C., and Heimpel, G. E. 2005. Impact of flowering buckwheat on Lepidopteran cabbage pests and their parasitoids at two spatial scales. Biological Control 34:290-301.

Legault, S., Hébert, C., Blais, J., Berthiaume, R., Bauce, E., & Brodeur, J. 2012. Seasonal ecology and thermal constraints of *Telenomus* spp. (Hymenoptera: Scelionidae), egg parasitoids of the hemlock looper (Lepidoptera: Geometridae). Environmental entomology 41:1290-1301.

Leius, K. 1967. Influence of wild flowers on parasitism of tent caterpillar and codling moth. The Canadian Entomologist 99:444-446.

Linit, M. J., and Stephen, F. M. 1983. Parasite and predator component of within-tree southern pine beetle (Coleoptera: Scolytidae) mortality. The Canadian Entomologist 115: 679-688.

Liu, C. L. 1926. On Some Factors of Natural Control of the Eastern Tent Caterpillar (*Malacosoma americana* Harris): With Notes on the Biology of the Host. Cornell University.

Lyons, L. A. 1962. The effect of aggregation on egg and larval survival in *Neodiprion swainei* Midd. (Hymenoptera: Diprionidae). The Canadian Entomologist 94:49-58.

Lyons, L. A. 1964. The European pine sawfly, *Neodiprion sertifer* (Geoff.) (Hymenoptera: Diprionidae). A review with emphasis of studies in Ontario. Proceedings of the Entomological Society of Ontario 94:5-37.

Marsh, P. M. 1968. The Nearctic Doryctinae, VII. The genus *Doryctes* Haliday (Hymenoptera: Braconidae). Transactions of the American Entomological Society 94:379-405.

Mason, W. R. M. 1978. A synopsis of the Nearctic Braconini, with revisions of Nearctic species of *Coeloides* and *Myosoma* (Hymenoptera: Braconidae). The Canadian Entomologist 110:721-765.

Mason, W. R. M. 1979. A new *Rogas* (Hymenoptera: Braconidae) parasite of tent caterpillars (*Malacosoma* spp. Lepidoptera: Lasiocampidae) in Canada. Canadian Entomologist 111: 783-786.

Mathews, P. L., and Stephen, F. M. 1997. Effect of artificial diet on longevity of adult parasitoids of *Dendroctonus frontalis* (Coleoptera: Scolytidae). Environmental entomology 26:961-965.

McGovern, W. L., Cross, W. H., and Mitchell, H. C. 1974. *Eupelmus cyaniceps* (Hymenoptera: Eupelmidae) a hyperparasite. Journal of the Georgia Entomological Society 9:68-69.

McGregor, M. D., and Sandin, L. O. 1968. Observations on the pinyon pine sawfly, *Neodiprion edulicolus*, in eastern Nevada (Hymenoptera: Diprionidae). The Canadian Entomologist 100:51-57.

Minnoch, M. W., and Parker, D. L. 1971. Life History of a Looper, *Lambdina punctata*, in Utah (Lepidoptera: Geometridae). Annals of the Entomological Society of America 64:386-389.

Morris, C. L., and Schroeder, W. J. 1966. Pine cone mortality. Va Forests 21:18, 20.

Moser, J. C., Thatcher, R. C., and Pickard, L. S. 1971. Relative Abundance of Southern Pine Beetle 1 Associates in East Texas. Annals of the Entomological Society of America 64:72-77.

Muesebeck, C. F. W. 1980. The Nearctic parasitic wasps of the genera *Psilus* Panzer and *Coptera* Say (Hymenoptera, Proctotrupoidea, Diapriidae). United States Department of Agriculture Technical Bulletin 1617:1-71.

Newcomer, E. J. 1936. Effect of cold storage on eggs and young larvae of codling moth. Journal of Economic Entomology 29:1123-1125.

Noyes, J. S. 2017. Universal Chalcidoidea Database. World Wide Web electronic publication. <http://www.nhm.ac.uk/chalcidoids>

Overgaard, N. A. 1968. Insects associated with the southern pine beetle in Texas, Louisiana, and Mississippi. Journal of Economic Entomology 61: 1197-1201.

Packard, A. S. 1881. Insects injurious to forest and shade trees. United States Entomological Commission Bulletin 7: 1-275.

Parry, D. 1995. Larval and pupal parasitism of the forest tent caterpillar, *Malacosoma disstria* Hübner (Lepidoptera: Lasiocampidae), in Alberta, Canada. The Canadian Entomologist 127:877-893.

Peck, O. 1963. A catalogue of the nearctic Chalcidoidea (Insecta: Hymenoptera). The Memoirs of the Entomological Society of Canada 95(S30):5-1092.

Porter, B. A. 1917. The host of *Ablerus clisiocampae* Ash. Entomological News 28:186.

Porter, B. A., and C. H. Alden. 1921. *Anaphoidea conotracheli* Girault (Hym.), and egg parasite of the apple maggot. Proceedings of the Entomological Society of Washington 23:62–63.

Price, P. W. 1970. Characteristic permitting coexistence among parasitoids of a sawfly in Quebec. Ecology 51:445-454.

Price, P. W., and Tripp, H. A. 1972. Activity patterns of parasitoids on the Swaine jack pine sawfly, *Neodiprion swainei* (Hymenoptera: Diprionidae), and parasitoid impact on the host. The Canadian Entomologist 104:1003-1016.

Raizenne, H. 1957. Forest sawflies of southern Ontario and their parasites. Canada Department of Agriculture Publication 1009:1-44.

Rauf, A., and Benjamin, D. M. 1980. The biology of the white pine sawfly, *Neodiprion pinetum* (Hymenoptera: Diprionidae) in Wisconsin. The Great Lakes Entomologist 13:219-224.

Riley, M. A., and Goyer, R. A. 1986. Impact of Beneficial Insects on *Ips* spp. (Coleoptera: Scolytidue) Bark Beetles in Felled Loblolly and Slash Pines in Louisiana. Environmental Entomology 15:1220-1224.

Saffer, B. 1982. Systematic revision of the genus *Cenocoelius* (Hymenoptera, Braconidae) in North America including Mexico. Polish Journal of Entomology 52:73-167.

Schaffner Jr, J. V. 1934. Introduced parasites of the brown-tail and gipsy moths reared from native hosts. Annals of the Entomological Society of America 27:585-592.

Shaw, M.R. 2002. Host ranges of *Aleiodes* species (Hymenoptera: Braconidae), and an evolutionary hypothesis. Pages 321-327 *in*: G. Melika and C. Thuróczy (eds.), Parasitic wasps: Evolution, Systematics, Biodiversity and Biological Control. Agroinforum, Budapest.

Shaw, S. R. 2006. *Aleiodes* wasps of eastern forests: a guide to parasitoids and associated mummified caterpillars. US Department of Agriculture, Forest Service.

Smith, D. R. 1996. Aulacidae (Hymenoptera) in the mid-Atlantic states, with a key to species of eastern North America. Proceedings of the Entomological Society of Washington 98:274-291.

Stacey, L., Roe, R., and Williams, K. 1975. Mortality of eggs and pharate larvae of the eastern tent caterpillar, *Malacosoma americana* (F.) (Lepidoptera: Lasiocampidae). Journal of the Kansas Entomological Society 48:521–523

Stehr, F. W., and E. F. Cook. 1968. Revision of the genus *Malacosoma* Hübner in North America (Lepidoptera: Lasiocampidae): systematics, biology, immatures, and parasites. Bulletin (United States National Museum) 276:1-321

Stelzer, M. J. 1968. The Great Basin tent caterpillar in New Mexico: life history, parasites, disease and defoliation. U.S. Forest Service, Rocky Mountain Forest Experimental Station Paper 39.

Stevenson, R. E. 1967. Notes of the Biology of the Engelmann Spruce Weevil, *Pissodes engelmanni* (Curculionidae: Coleoptera) and its Parasites and Predators. The Canadian Entomologist 99:201-213.

Struble, G. R. 1957. Biology and control of the white-fir sawfly. Forest Science 3:306-313.

Sullivan, B. T., Pettersson, E. M., Seltmann, K. C., and Berisford, C. W. 2000. Attraction of the bark beetle parasitoid *Roptrocerus xylophagorum* (Hymenoptera: Pteromalidae) to host-associated olfactory cues. Environmental Entomology 29:1138-1151.

Townes, H., and Townes, M. 1960. Ichneumon-Flies of America North of Mexico Pt. 2: Subfamilies Ephialtinae, Xoridinae, and Acaenitinae. Memoirs of the American Entomological Institute 12:1-537.

Treherne, R. C. 1921. A further review of applied entomology in British Columbia. Journal of the Entomological Society of British Columbia 135-146.

Vanlaerhoven, S. L., Stephen, F. M., and Browne, L. E. 2005. Adult parasitoids of the southern pine beetle, *Dendroctonus frontalis* Zimmermann (Coleoptera: Scolytidae), feed on artificial diet on pine boles, pine canopy foliage and understory hardwood foliage. Biocontrol science and technology 15:243-254.

Walkley, L. M. 1954. A new cryptine genus of economic interest (Hymenoptera: Ichneumonidae). Journal of the Washington Academy of Sciences 44:219-220.

Wegensteiner, R., Wermelinger, B., and Herrmann, M. 2015. Natural enemies of bark beetles: predators, parasitoids, pathogens, and nematodes. Pages 247-304 i*n* F. E. Vega & R. W. Hofstetter (eds.), Bark Beetles. Biology and Ecology of Native and Invasive Species Elsevier: Amsterdam

Wellington, W. G. 1965. Some maternal influences on progeny quality in the western tent caterpillar, *Malacosoma pluviale* (Dyar). The Canadian Entomologist 97:1-14.

Wetzel, B. W., Kulman, H. M., and Witter, J. A. 1973. Effects of Cold Temperatures on Hatching of the Forest Test Caterpillar, *Malacosoma disstria* (Lepidoptera: Lasiocampidae). The Canadian Entomologist, 105:1145-1149.

Wharton, R.A., 1997. Generic relationships of opiine Braconidae (Hymenoptera) parasitic on fruit-infesting Tephritidae (Diptera). Journal of the Washington Academy of Sciences. 68:147–167.

Wharton, R. A. and Marsh, P. M. 1978. New World Opiinae (Hymenoptera: Braconidae) parasitic on Tephritidae (Diptera). Journal of the Washington Academy of Sciences 68:147-167.

Wharton, R., Ward, L., and Miko, I. 2012. New neotropical species of Opiinae (Hymenoptera, Braconidae) reared from fruit-infesting and leafmining Tephritidae (Diptera) with comments on the *Diachasmimorpha mexicana* species group and the genera *Lorenzopius* and *Tubiformopius*. ZooKeys 243:27-82.

Wharton, R. A., and Yoder, M. J. Parasitoids of Fruit-Infesting Tephritidae. http://paroffit.org. Accessed on Wed Dec 06 12:22:40 -0600 2017.

Wilkinson, R. C. 1971. *Neodiprion excitans* (Hymenoptera: Diprionidae) on sand pine in Florida. The Florida Entomologist 54:343-344.

Wilkinson, R. C., Becker, G. C., and Benajmin, D. M. 1966. The biology of *Neodiprion rugifrons* (Hymenoptera: Diprionidae), a sawfly infesting jack pine in Wisconsin. Annals of the Entomological Society of America 59:786-792.

Wilkinson, R. C., and Chellman, C. W. 1978. A new sawfly on sand pine in West Florida. Florida entomologist 61:26.

Wilkinson, R. C., and Drooz, A. T. 1979. Oviposition, Fecundity, and Parasites of *Neodiprion excitans* from Belize, CA. Environmental Entomology 8:501-505.

Williams, L. T. 1916. Notes on the egg-parasites of the apple tree tent-caterpillar (*Malacosoma americanum*). Psyche 23:148-153.

Witter, J. A., and Kulman, H. M. 1972. Review of the parasites and predators of tent caterpillars (*Malacosoma* spp.) in North America. Agricultural Experiment Station University of Minnesota. Techincal Bulletin 289.

Witter, J.A. and Kulman, H.M., 1979. The parasite complex of the forest tent caterpillar in northern Minnesota. Environmental Entomology, 8:723-731.

Yee, W. L., Goughnour, R. B., Hood, G. R., Forbes, A. A., and Feder, J. L. 2015. Chilling and host plant/site-associated eclosion times of Western cherry fruit fly (Diptera: Tephritidae) and a host-specific parasitoid. Environmental Entomology 44:1029-1042.
